# Farmer’s perspectives of rodent pests’ damage and management practices in Wenchi highlands, central Ethiopia

**DOI:** 10.1101/2022.12.06.519363

**Authors:** Kabeta Legese, Afework Bekele

## Abstract

A study was conducted to obtain information about the knowledge, attitudes and practices (KAP) of farmers in Wenchi highlands on rodent damage and their management practices. Farmers (n=395) from four highland villages of Wenchi District were randomly selected and interviewed using a semi-structured questionnaire. Rodents were identified as major pests, and perceived negatively among farmers. There were significant variations in the type of damage (χ2=112.698, df= 3, p < 0.05) and crop types susceptibility to rodent pest attack (χ2= 143.26, df = 3, P < 0.05). Crop damage (38.7%) and damage to human properties (27.9%) were the two dominant rodent related problems in the area. Barley was the most susceptible crop to rodent attack (57.5%). The occurrence of frequency of rodent pests and crop damage between the cropping stages also varied significantly. Most damage on barley crop (42.5%) occurred during the maturation stage. Farmers assessed and detected rodents damage by observing damaged seeds, damaged stores and rodent droppings in the storage, and stem cut of standing crops in the crop fields. None of the farmers have employed any management strategy in barley crop fields stating that this is practically impossible. In storage, farmers mainly use cats (53.73%) and trapping (22.64%) to control rodents. Detailed off-field rodent damage assessment, and community education for rodent management are recommended.

## 1. Introduction

Rodents have played an important part in human history as a food source (Palis *et al*., 2011), model animals for research and good indicators of environmental quality (Habtamu and Bekele, 2012). They are also a threat to food production and human property, and are a public health risk (Panti-May *et al*., 2017). Consequently, rodents are considered as the most serious vertebrate pests worldwide (Staples *et al*., 2003; Brown *et al*., 2007). However, only <10% of rodent species are major pest species, and even fewer cause problems in broader areas (Stenseth *et al*., 2003; Capizzi *et al*., 2014).

Rodent pests remarkably affect the global crop production and livelihoods of farmers because their cost to agriculture is enormous (Singleton *et al*., 2005, 2010). They cause huge economic losses in agricultural crops, mainly root crops and cereals in the field, and consume and contaminate stored grains (Mulungu *et al*., 2005; Yonas *et al*., 2010; Mulungu *et al*., 2015; Jones *et al*., 2017). Rodents consume foodstuffs, cause physical damage to packaging and storage materials, and contaminate products with hair, urine and feces (Brown *et al*., 2013; Buckle, 2015; Lund, 2015). They are responsible for damaging food volume that could feed about 280 million people for a year (Meerburg *et al*., 2009). Rodent hair or droppings in food may create great problems for exporting countries up to the rejection of the entire loads (Lund, 2015).

Rodent damages significantly affect food security and income of small-holder farmers in developing countries (Brown *et al*., 2013; Lund, 2015; Htwe *et al*., 2016). The damage can be severe, diverse, and show temporal and spatial variations (Gebhardt *et al*., 2011) because it is directly associated with rodent abundance, diversity, feeding habits and reproductive patterns (Swanepoel *et al*., 2017). Crop losses also vary between crops, cropping stages, and storage types (Swanepoel *et al*., 2017). Annual losses due to rodents in several countries are economically unacceptable (Yonas *et al*., 2010). Such a country level damage can have a major effect on the economy of any country and all available consumers (Gebhardt *et al*., 2011).

Rodents are one of the major problems in Africa, and have been the number one crop pest (Hill, 2008). Ethiopia also experiences constant rodent pest problems on different agricultural crops (Bekele *et al*., 2003; Makundi *et al*., 2005). Bekele *et al*. (2003) have recorded the highest rodent crop damages (26%) in central Ethiopia. A more recent report has revealed that Ethiopia is the third country in stored grain losses after Egypt and Tanzania (Mulungu, 2017).

It is economically beneficial to control rodent population to reduce rodent linked losses (Skonhoft *et al*., 2006; Brown *et al*., 2013). This is influenced by the farmer’s knowledge on variables affecting crop damage, the level of crop susceptibility, the rodent pest population during the most susceptible crop stage and how much they are prepared to control the pests (Makundi *et al*., 2005). Local perceptions about rodents and the damage they cause are vital, as a first step, to design and implement rodent control or educational programs (Garba *et al*., 2013; Panti-May *et al*., 2017). Studies have been carried out on this subject in northern and southern Ethiopia (Yonas *et al*., 2010; Tomass *et al*., 2020), but still not documented from Wenchi highlands. Thus, the purpose of this survey was to assess the knowledge, attitudes and practices (KAP) of farmers on rodent damage and management in Wenchi highlands.

## 2. Methods

### 2.1. Study area description

The study was conducted in the central highlands of Ethiopia, Wenchi district of southwest Shewa Zone, Oromia. It is located between Ambo and Waliso towns, 155 km away from the capital, Addis Ababa (Fig. 1). The altitude of the area ranges between 2,810 and 3,386 m above sea level (Tefera *et al*., 2002). Its highest elevation is at Mount Wenchi (3,386 m asl). The area is characterized by highland sub-humid climate with the average annual rainfall of 1400 to 1420 mm (Shale *et al*., 2014; Angessa *et al*., 2020). The area receives unimodal rainfall with longer rainy periods stretching from May to September. The peak rainfall occurs in July and August. The cold-dry season is distinguished between October and January (Degefu and Schargel, 2015; Angessa *et al*., 2020). The temperature varies from 14 to 26°C during the day and falls below 10°C at night (Degefu and Schargel, 2015).

**Figure 1.**
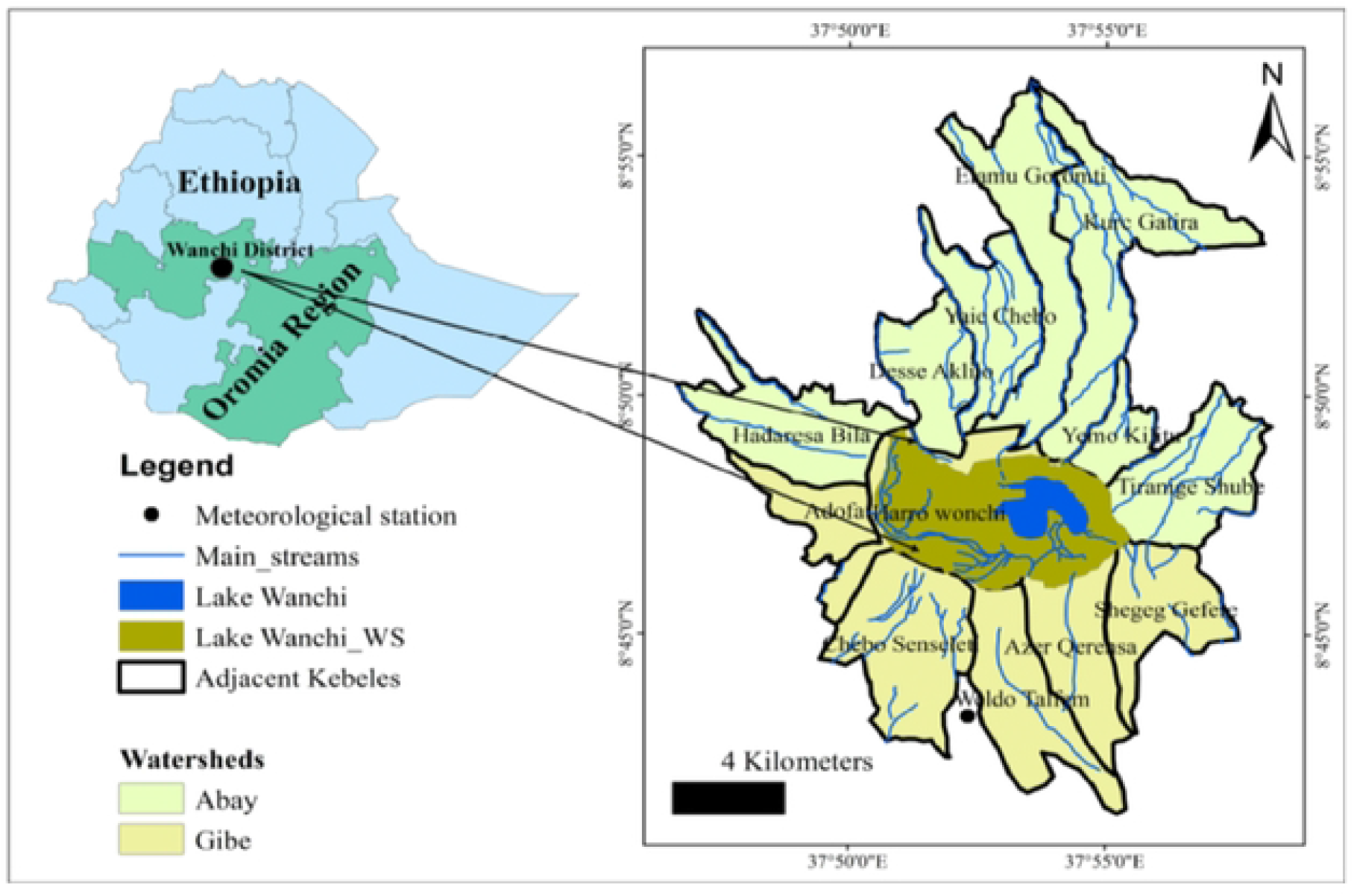
Map of the study area (Adopted from Angessa *et al*., 2022)

Lake Wenchi is among a few remaining fairly pristine high-mountain crater lakes in the central highlands (Degefu and Schagerl, 2015; Angessa *et al*., 2020). It is one of the popular tourist attractions and interesting ecotourism destinations in this area (Fig. 1). As a result, Oromia Tourism Commission has recognized it as the best national tourism destination area in the region. The area also owns a 15th century Monastery and a hilly highland area covered with natural forests, mineral waters and hot springs (Shale *et al*., 2014; Degefu *et al*., 2014).

The main livelihood in the area is mixed agriculture (crop cultivation and livestock rearing), small and micro-enterprises, and income generating activities from ecotourism (Shale *et al*., 2014; Angessa *et al*., 2022). The average land holding size for a single household is 0.5 hectares and the major crops grown in the area are enset (*Ensete ventricosum*), barely, wheat, and potato (Shale *et al*., 2014). Like other highland farmers of the country, farming and harvest is performed by traditional technology (Yonas *et al*., 2010).

### 2.2. Farmer surveys

A total of four relatively accessible highland villages of Wenchi District were purposely selected in reference to Lake Wenchi and Haro town. These villages were also located either in or adjacent to the area where the ongoing rodent ecology research project is being conducted by the same group of researchers. The questionnaires were administered household farmers randomly selected from lists obtained from the administration bureau of the respective villages. Household samples were computed using the estimation formula for a single proportion, *n* = *Z*^2^2*P*(1 − *P*)/*d*^2^; *w*here *n* is calculated sample size, *Z* is critical value (1.96) at 95 % confidence level, *P* is an expected proportion (50%) and *d* is precision or margin of error which is fixed at 5% (Daniel, 1995).

Moreover, the calculated sample size was proportionally assigned to each of the four villages using the formula, *nh* = *N*(*n*)/∑N (Pandey and Verma, 2008); where, *nh* is the number of sample households to be selected from each village **N** is the total number of households in each village *n* is the calculated total sample size to be selected from all the study villages, and ∑N is the sum total of households in the selected villages. Thus, the questionnaires were administered to a total of 395 randomly selected households from lists obtained from the administrations of the respective villages (Table 1). Individual household was randomly selected from the study villages by lottery method for data collection.

**Table 1.** The distribution of sampled households in the study villages

| Villages | No. of households |  |  | No. of sampled households |  |  |
| --- | --- | --- | --- | --- | --- | --- |
|  | Male headed | Female headed | Total | Male headed | Female headed | Total |
| Haro Wenchi | 746 | 106 | 856 | 91 | 12 | 103 |
| Waldo Telfami | 595 | 184 | 883 | 84 | 22 | 106 |
| Cabo Sansalati | 573 | 108 | 681 | 69 | 13 | 82 |
| Azar Qeransa | 746 | 114 | 860 | 91 | 13 | 104 |
| Total | 2660 | 512 | 3172 | 335 | 60 | 395 |

A semi-structured questionnaire was extracted from published studies with similar objectives (Brown *et al*., 2008; Yonas *et al*., 2010; Stuart *et al*., 2011; Tomass *et al*., 2020) and modified to the situation of the area. It was designed not to be too troublesome and long for farmers, while enabling the collection of information relevant to the research questions (Foster *et al*., 2019). Both open and closed ended questions were prepared in English and administered using local language (Afan Oromo) to one person to provide accurate information (Garba *et al*., 2013). Each interview was conducted by one of the researchers and his field assistant for approximately 30 minutes between in December, 2020 and February, 2021.

The questionnaire was composed of four parts. In the first part, demographic profiles, such as age and level of education of the respondents were collected. The second section contained questions that tested the agricultural practices in the respective villages. The third section gathered information on farmers’ knowledge of rodent damages and their perceptions. In the fourth and last section, rodent management practice data were obtained.

### 2.3. Data analyses

The collected data were coded, cleaned and summarized using Microsoft Excel spread sheet. Both qualitative and quantitative data were analyzed with appropriate statistical methods such as mean, percentage, and Chi-square test using SPSS version 20 (SPSS, Inc. USA). Chi-square (χ2) tests were used to verify possible associations between socioeconomic profiles of the respondents and their response to KAP questionnaires. The differences between the villages in farming practices and composition were also compared using Chi-squared tests. Probability values were considered statistically significant when P-value is ≤ 0.05.

## 3. Results

### 3.1. Socio–demographic characteristics of farmers

Out of the 395 study participants, 335 (84.8%) were males and 60 (15.2) females. The majority of the respondents (82.2%) were between 20 and 50 age ranges. Nearly half (47.8%) of the study participants were not registered for any formal education, while 36.9% attended a primary education. Majority of the respondents (86.5%) had between 5 and 10 family sizes, while the remaining had above 3 family members. The respondents have spent from 5 years to more than 50 years on farming. Most of the respondents (78.2%) had a farmland size below one to two hectares, while only 7. 3% of them had more than three hectares. Most (84.8%) of the respondents supported their livelihoods through mixed farming practices. Only 15.2% of the respondents generated additional income through ecotourism (Table 2). Rodent crop damage and type of management were not significantly explained by village, age, family size, and education.

**Table 2.** Socio–demographic characteristics of the study participants

| Variables |  | No. of respondents | Percentage (%) |
| --- | --- | --- | --- |
| Sex | Male | 335 | 84.8 |
|  | Female | 60 | 15.2 |
| Age in years | 20–30 | 103 | 26.07 |
|  | 31–40 | 138 | 34.93 |
|  | 41–50 | 84 | 21.26 |
|  | >50 | 70 | 17.72 |
| Educational status | Illiterate | 189 | 47.84 |
|  | Primary education | 146 | 36.96 |
|  | Secondary education | 23 | 15.18 |
| Family size | <5 | 117 | 29.62 |
|  | 5–10 | 225 | 56.96 |
|  | >10 | 53 | 13.41 |
| Farming years | <10 | 49 | 12.4 |
|  | 10–20 | 90 | 22.78 |
|  | >20 | 256 | 64.81 |
| Farmland size | <1ha | 215 | 54.43 |
|  | 1–2ha | 94 | 23.79 |
|  | 2.1–3ha | 57 | 14.43 |
|  | >3ha | 29 | 7.34 |
| Economic activity | Ecotourism | 60 | 15.18 |
|  | Mixed farming | 335 | 84.82 |

### 3.2. Rodent pests and crop damages

Crop and livestock productions are the main sources of income for the farmers in Wenchi highlands. The major crop types grown in the study area are barley, wheat, enset and potato. Barley and enset were the two chief crop types produced by all the farmers. In addition to these to crops, wheat and potato were produced in relatively lower and upland areas, respectively. There was insignificant difference in the responses of farmers on the major crops grown in the area (χ2=4.725, df=3, P>0.05).

Results from farmers’ interviews and indirect observations revealed that rodents including mole rats and porcupines were the major crop pests in the area. *Arvicanthis abyssinicus, Mastomys natalensis, M. awashensis, Hystrix cristata* and *Tachyoryctes splendens* were the five rodent pest species recorded from their occurrences in the neighboring natural habitats, indirect evidences and the surrounding community reports. The first three pests were live and snap-trapped from the adjoining forest remnants, while the remaining two were documented through indirect evidence and reports from the surrounding community.

The current study showed that rodents were the most important pest to farmers. The farmers also believed that rodents are useless, and a damaging nature. The farmers had cited crop damage, disturbance, food contamination and damage to human properties as the most rodent inflicted problems. Crop damage (38.7%) and damage to human properties (27.9%) were the two predominant rodent related problems in the area (Table 3). There was a significant difference in types of rodent damage in the study area (χ2=112.698, df= 3, p < 0.05).

**Table 3.**
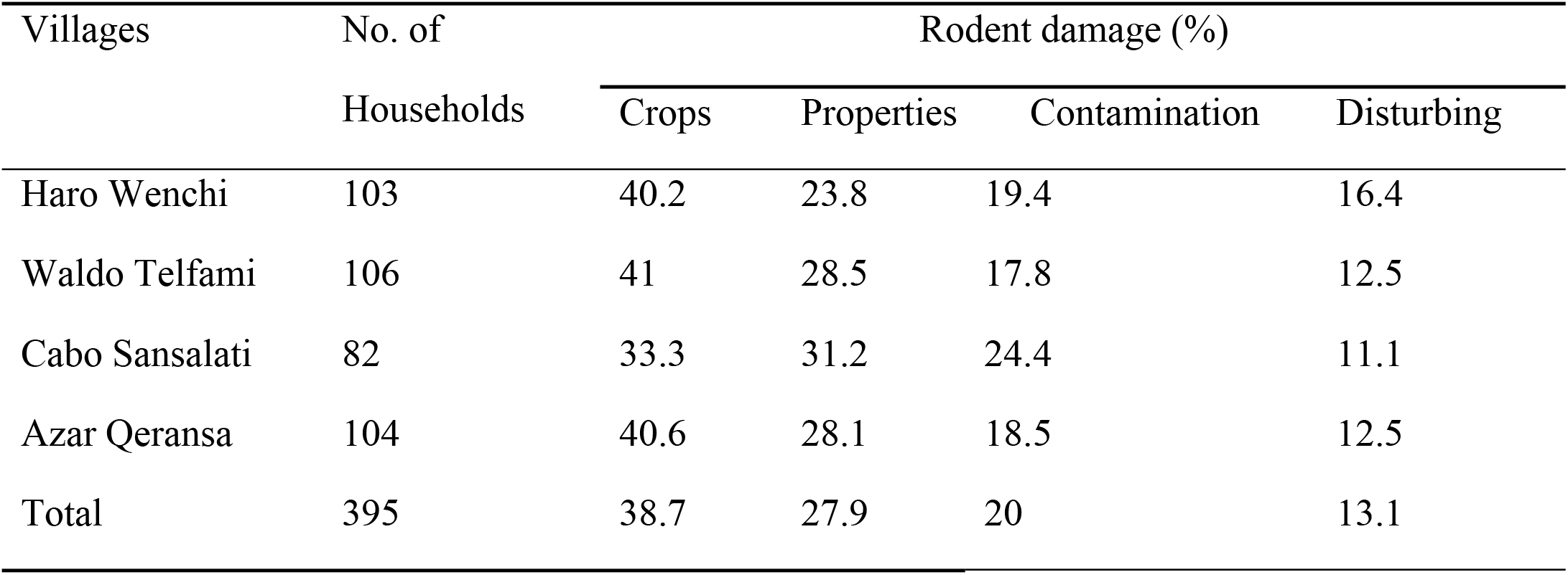
Types of rodent damage in Wenchi highlands

| Villages | No. of Households | Rodent damage (%) |  |  |  |
| --- | --- | --- | --- | --- | --- |
|  |  | Crops | Properties | Contamination | Disturbing |
| Haro Wenchi | 103 | 40.2 | 23.8 | 19.4 | 16.4 |
| Waldo Telfami | 106 | 41 | 28.5 | 17.8 | 12.5 |
| Cabo Sansalati | 82 | 33.3 | 31.2 | 24.4 | 11.1 |
| Azar Qeransa | 104 | 40.6 | 28.1 | 18.5 | 12.5 |
| Total | 395 | 38.7 | 27.9 | 20 | 13.1 |

Most of the respondents claimed the vulnerability of all the crops grown in the area to rodent damage. However, barley (57.5%) was reported as the most affected crop by rodents followed by root crops (Table 4). Mice and rats damage barley and wheat, while mole rats and porcupines damage enset and potato. There was a significant variation among the major crop types susceptibility to rodent pest attack (χ2= 143.26, df = 3, P < 0.05).

**Table 4.** Types of crops grown and their susceptibility to rodent attack

| Villages | No. of household | Damage due to rodent pests (%) |  |  |  |
| --- | --- | --- | --- | --- | --- |
|  |  | Barley | Enset | Potato | Wheat |
| Haro Wenchi | 103 | 52.23 | 19.40 | 26.86 | 4.47 |
| Cabo Sansalati | 106 | 48.21 | 16.07 | 23.21 | 12.5 |
| Waldo Telfami | 82 | 51.11 | 33.33 | 26.66 | 11.11 |
| Azar Qeransa | 104 | 43.75 | 18.75 | 34.37 | 6.25 |
| Total | 395 | 57.5 | 13.5 | 23.0 | 6.0 |

The annual crops yield varied among the farmers in the study area in relation to their farmland size. Most of the farmers (54.5%) obtain very low amount of crop yield, which is less than 5 quintals in average. Only few the farmers (21.7%) harvest more than 15 quintals average yields (Fig. 2). There was a statistically significant differences in the annual crop yields among farmers (χ2= 214. 451, df= 4, P <0.05).

**Figure 2.**
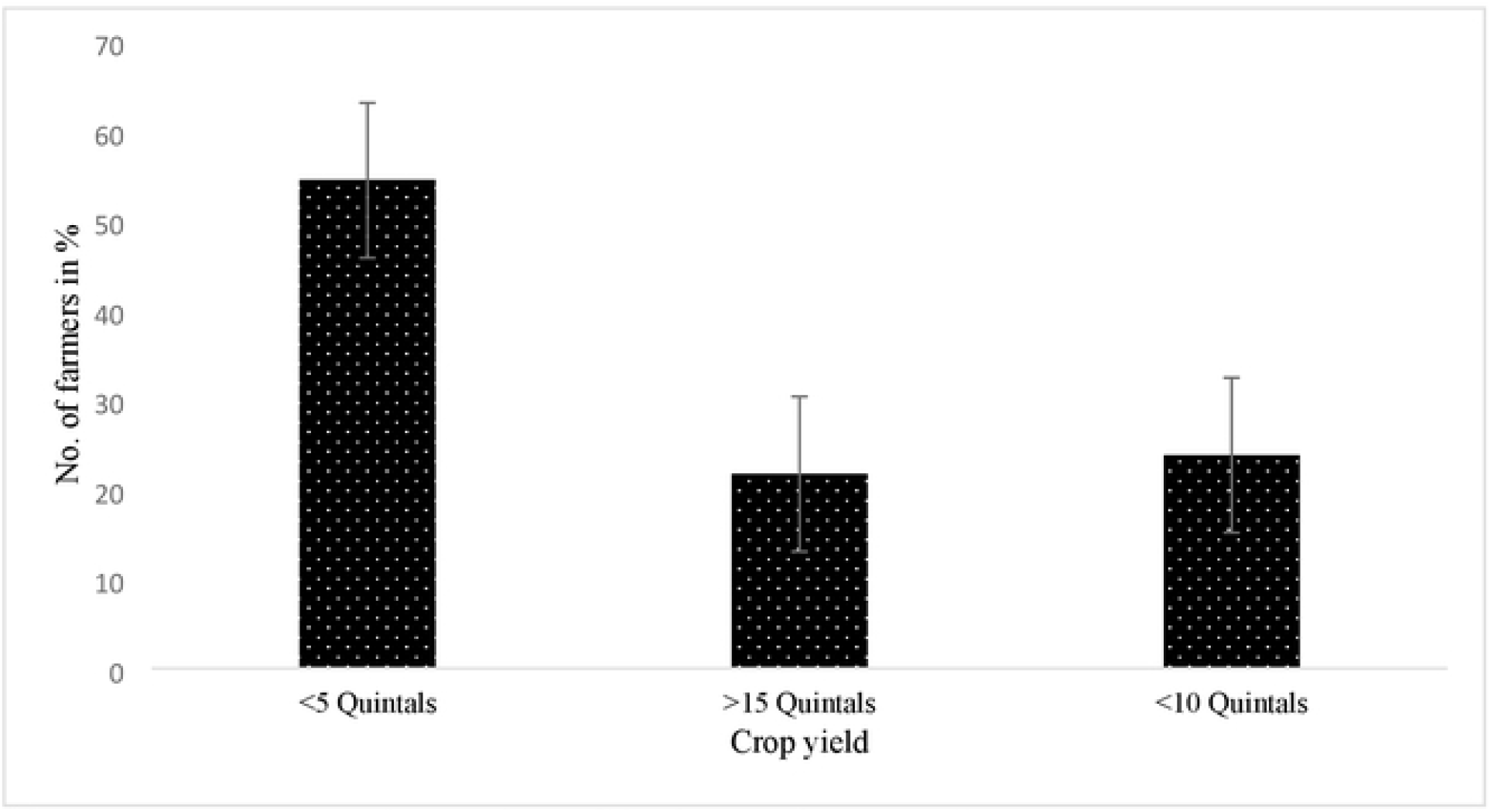
Average annual crop yield of the farmers in the study area

Farmers also crudely estimated crop losses due to rodent pests. Most respondents (74.5%) estimated an average of 1.5 quintals of crops might be damaged by rodents in the storage. However, they were unable to estimate crop damage in the crop fields in figures. The level of rodent crop damage in the area is generally high. Most of the respondents (87. 4%) associated a high crop damage to rodent pests in the area. Only less than 10% of the farmers reported low crop damage by rodent pests (Fig. 3). There was a statistically significant variation in the responses of farmers on the level of crop damage by rodents (χ2= 196. 371, df= 2, P < 0.05).

**Figure 3.**
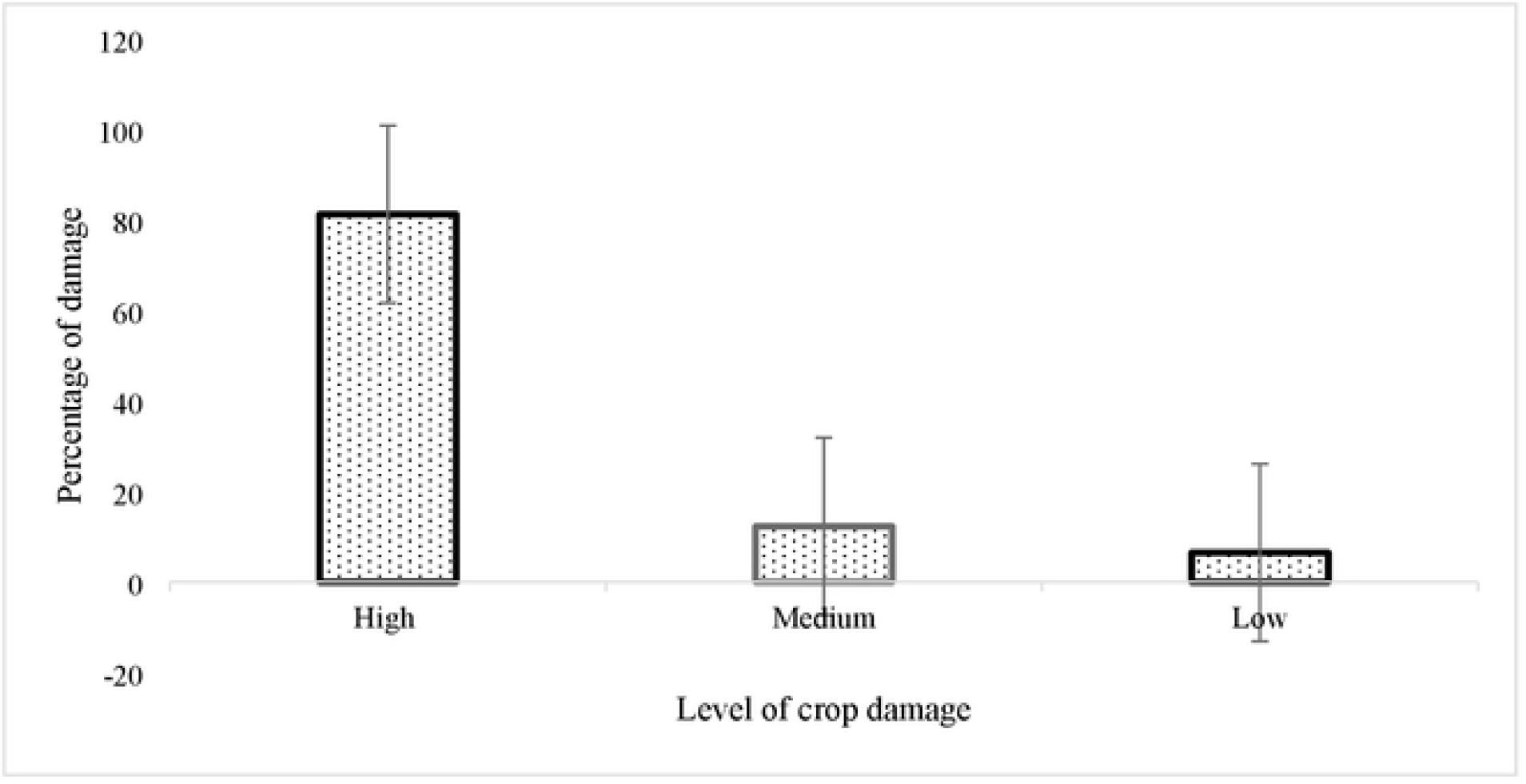
Level of crop damage by rodent pests in the study area

The majority of farmers (72.5%) reported a seasonal variation of rodent damaging behavior in the house as well as the crop field. House rodent infestation was higher during the wet seasons, but the damage was significant during the dry season in the crop fields. Most farmers reported regular presence of rodent damage (in every cropping season/year) in their locality (Table 5). The responses of farmers on the occurrence frequency of rodent pests varied significantly (χ2=193.826, df = 2, P < 0.05).

**Table 5.** Frequency of rodent crop damage in Wenchi highlands

| Villages | No. of households | Frequency of crop damage (%) |  |  |
| --- | --- | --- | --- | --- |
|  |  | Regular | Frequent | Occasional |
| Haro Wenchi | 103 | 56.71 | 25.37 | 17.1 |
| Waldo Telfami | 106 | 41.07 | 26.78 | 32.14 |
| Cabo Sansalati | 82 | 44.44 | 24.44 | 33.33 |
| Azar Qeransa | 104 | 43.75 | 21.87 | 32.37 |
| Total | 395 | 47.5 | 25.0 | 28.0 |

Crop damage by rodent pests occurred during both post-harvest and pre-harvest stages. Farmers identified rodent damage at different cropping stages starting from sowing to harvesting for different crop types. They have noted serious damage on barley (42.5%) and enset (35%) crops during maturity. These crops were also vulnerable to rodent damage during vegetative and booting stages. Potatoes were highly damaged both during sowing and after their maturity (Fig. 4). There was a significant difference in crop damage between the cropping stages (χ2= 110.82, df = 2, P < 0.05).

**Figure 4.**
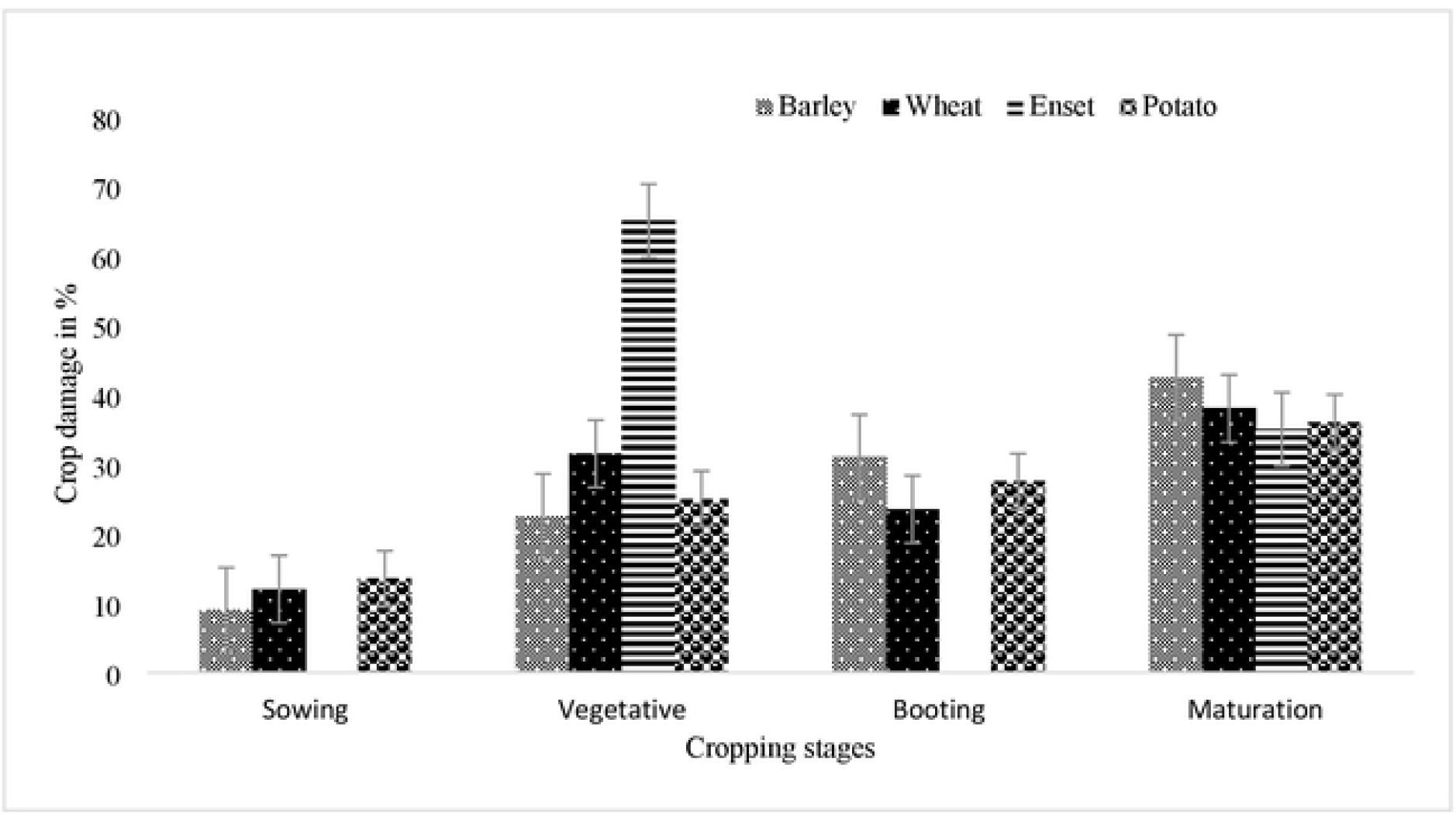
Rodent crop damage in different cropping stages in the study area

Most farmers in the study area supervise their farms occasionally before harvest and rarely after harvesting. There was no significant variation on farm supervision practices among farmers (χ2=1.691, df = 1, P > 0.05). Farmers assess and detect the presence and damage of rodents in the storage and crop fields using different assessment mechanisms. Observation of damaged seeds (32%), damaged stores (27%) and rodent droppings (23%) were the most used methods to detect the damage or/and presence of rodents in the house. Stem cut of standing crops (73.5%) was the most used assessment method of rodent damages in the crop fields (Fig. 5).

**Figure 5.**
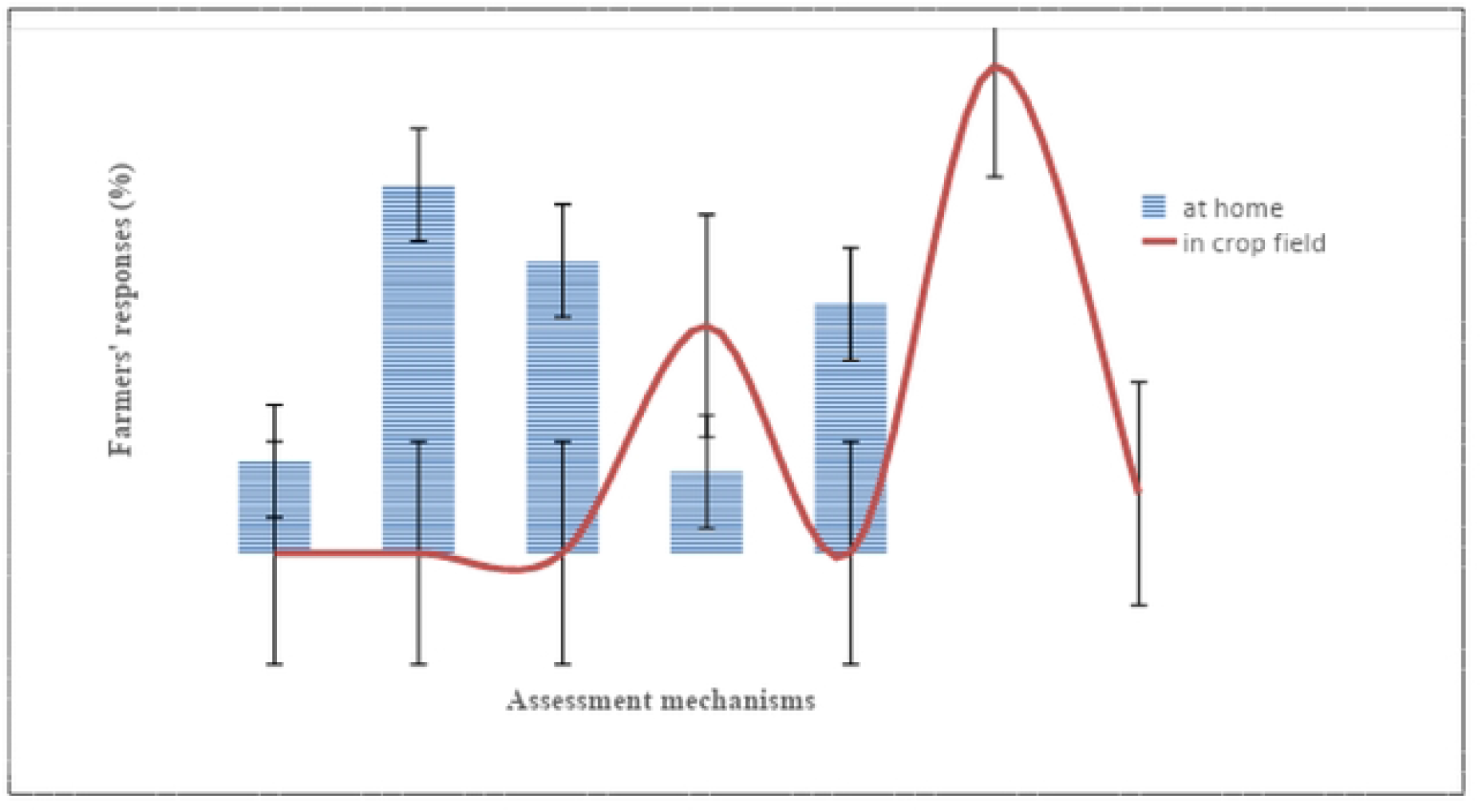
Farmer’s assessment mechanisms of rodent damage in the house and crop fields

### 3.3. Rodent management

The farmers in the study area employ different management methods to control rodent damage in storage (Table 6). The farmers used domestic cats (53.73%) followed by trappings (22.64%) to contain rodent damage in storage. A significant variation was shown among the rodent pest preventing mechanisms used by the respondents during storage (χ2= 89.63, df= 3, P < 0.05). Trapping and hunting were employed to control rodent damage on enset and potatoes. However, none of the interviewed farmers have employed any management strategies in the barley crop fields. Farmers in the study area claimed that rodenticides were not safe, and rodents have developed an adaptation to avoid rodenticides and traps.

**Table 6.** Rodent pest management techniques used in the study area during storage

| Villages | No. of households | Management strategies (%) |  |  |  |
| --- | --- | --- | --- | --- | --- |
|  |  | Cats | Trapping | Rodenticide | Hunting |
| Haro Wenchi | 103 | 53.73 | 28.35 | 7.46 | 10.44 |
| Waldo Telfami | 106 | 37.5 | 28.57 | 12.5 | 21.42 |
| Cabo Sansalati | 82 | 40.0 | 22.22 | 28.89 | 8.89 |
| Azar Qeransa | 104 | 34.37 | 25.0 | 21.87 | 18.75 |
| Total | 395 | 43.0 | 22.0 | 16.0 | 14.5 |

## 4. Discussion

The farmers in Wenchi highlands totally rely on farming and rearing cattle for their livelihoods. Only a few of them have generated additional income from ecotourism as documented by Shale *et al*. (2014) and Angessa *et al*. (2020). Six rodent species were identified as major crop pests in the area. Similarly, different rodent pest species were reported from numerous localities of Ethiopia (Bekele *et al*., 2003; Makundi *et al*., 2005; Yonas *et al*., 2010; Kasso, 2013; Tomass *et al*., 2020).

The current study showed that rodents damage the commonly grown crops, and are the most important pests in the area. This finding conforms with the worldwide problem associated with rodents (Staples *et al*., 2003; Brown *et al*.,2007). The farmers were well aware of rodent problems and expressed their frustration and anger towards these mammals. This result is consistent with earlier finding from Ethiopia (Makundi *et al*., 2005; Meheretu *et al*., 2010) and elsewhere (Makundi *et al*., 2005; Brown and Khamphoukeo, 2010). Furthermore, the farmers believe that rodents are useless but only damaging creatures. Such perception was common among sexes, all age groups, and area inhabitants. This finding is also in agreement with the findings from northern Ethiopia (Meheretu *et al*., 2014) and India (Singla *et al*., 2012). Local inhabitants were even viewing us as witchcraft when we were conducting field surveys. There were also occasions when individuals of different ages were harassing us verbally and intimidating us physically only because we had made contact with rodents. This implies that there is a big knowledge gap about the biological and ecological values of these natural biotas, and a need for a community wide education and training program on this matter.

In the present study, there was a significant difference in types of rodent damage in the study area. A closely related finding has been reported by Tomass *et al*. (2020) and Panti-May *et al*., (2017). Crop damage and damage to properties were the two dominant rodent attacks in the area. This is also in agreement with the finding of Panti-May *et al*. (2017). Like a report of Garba *et al*. (2013) from Niger, there is also an apparent absence of knowledge about the potential role of rodents in some public health issues.

Rodent crop damage varied between crops. This is in agreement with the finding of Swanepoel *et al*. (2017). Barley was the most affected crop by rodent pests followed by potatoes and enset. This is in agreement with the finding of Yonas *et al*. (2010) from the highlands of Tigray. However, it goes against several reports from the lowlands, where maize is the most susceptible crop to rodent attacks (Bekele *et al*., 2003; Makundi *et al*., 2005; Tomass *et al*., 2020). This difference is associated with variation in climatic conditions, the type of crop grown, and the wide distribution of these crops, and the inherent crop preferences of the present rodent pest species in these areas.

In the current study, farmers have observed rodent damage of barley from the sowing to harvesting stages. Similar results were reported by Yonas *et al*. (2010) and Wondifraw *et al*. (2021) from northern Ethiopia. However, the damage is higher in the maturation stage, when the barley is near to harvesting. This is in agreement with the experimental finding of Wondifraw *et al*. (2021) from south Gondar and the general patterns of rodent damage in the field crops (Lund, 2015; Swanepoel *et al*., 2017). The finding is, however, against a report from northern Ethiopia, where damage is severe at booting stages (Yonas *et al*., 2010). This variation might be associated with the difference in the accessibility and vulnerability of the crop, species richness and abundance of the rodent pests in the study areas.

The farmers reported a seasonal variation in rodent infestation of residential houses and crop fields. This is in congruent to the finding of Gebhardt *et al*. (2011) since damage is directly linked with rodent abundance, diversity, feeding habits and reproductive patterns (Swanepoel *et al*., 2017). However, it disagrees with a report of Staurt *et al*. (2011). This difference could be associated with the difference in geography, pest species, crop types and climatic variations between the areas.

The farmers have crudely estimated crop losses due to rodent pests. The estimation is in the same range of experimentally proved barley crop loss in south Gondar (Wondifraw *et al*., 2021). However, it is lower than the report from Tigray (Yonas *et al*., 2010) and southern Ethiopia (Tomass *et al*., 2020). This discrepancy might be due to the occasional supervision of crop fields by farmers, and the result confirms that rodent damage significantly affects food security and income of small-holder farmers of developing countries (Brown *et al*., 2013; Lund, 2015; Htwe *et al*., 2016). It also suggested the need of on-field rodent damage assessment to figure out the actual crop damage inflicted by rodents in the field.

In the present study, most farmers carried out farm supervision on rare occasions. The result is against a report of Yonas *et al*. (2010) from Tigray. This might be due to the belief that farmers are powerless to control the damage caused by rodents in the field in the study area. In consistent with reports from Tigray (Yonas *et al*., 2010) and Tanzania (Mulungu *et al*., 2015), rodent damage was assessed by observing damaged seeds, damaged stores and rodent droppings in the storage, and stem cut of standing crops crop fields.

Farmers in the study area have employed several indoor and outdoor rodent pest management strategies. Similar findings were reported similar practices by farmers (Yonas *et al*., 2010; Mulungu *et al*., 2015; Tomass *et al*., 2020). Farmers have used trapping and hunting to control rodent damages on enset and potatoes. These methods are well documented and the most practiced rodent control techniques in Ethiopia (Makundi *et al*., 2005). But it is against the finding of Yonas *et al*. (2010) where farmers were reliant on rodenticides for rodent pest management. This difference could be due to the farmers disregard of rodenticides by citing rodent adaptation and its side effects.

In the current study, none of the interviewed farmers have employed any management strategies in the barley crop fields. This is against the finding of Mulungu *et al*. (2015) from Central-eastern Tanzania, Tomass *et al*. (2020) from southern Ethiopia and Yonas *et al*. (2010) from northern Ethiopia, where most farmers used rodenticides in the crop fields. The farmers believe that managing rodents in the barley crop fields is practically not possible. This is in disagreement with the finding of Brown and Khamphoukeo (2010), but in total agreement with other studies conducted elsewhere in the world (Singleton and Petch, 1994; Palis *et al*., 2011; Singla *et al*., 2012). In these areas, many farmers accepted that they have little control over the damage to crops caused by rodents. Asian farmers, for instance, have a long history of planting two rows of grain for every 10 sown rows for rodents (Singleton and Petch, 1994). This might be due to the fact that rodents are minor and sporadic pests in the area, and are often ignored by farmers. A similar scenario has been reported from India (Singla *et al*., 2012).

Another possible reason that leads the farmers to a level of acceptance of rodent crops damage could be the chronic and prolonged nature of rodent depredation. This is the most likely rationalization for the current study area since it is experimentally supported, and Ethiopia experiences chronic rodent pest problems on different agricultural crops (Mulungu, 2017). This situation is unbearable in Ethiopia because it is experienced by small farm holders and the country is also facing an ever-increasing population and stunning economic inflation.

The farmers in the study area have claimed that rodents have developed an adaptation to avoid rodenticides and traps. However, these claims are unsubstantiated, and rodenticide avoidance of rodents could be associated with the quality of baiting foods. It is well documented that the use of poor baiting food has low rodent attraction potential to the rodenticide and leads to the point where rodents avoid consumption (Hill, 2008). Using high quality and palatable baits can easily reduce such problems. The second and most likely problem in the area could be the dose level of rodenticide preparation. This problem can be easily solved by balancing the amount of rodenticide and baiting food that is efficient and effective in attracting rodents. But, if rodents really developed rodenticide avoidance behavior, the only possible solution could be the use of other forms of rodenticides and rodent management strategies.

The claim of farmers in rodent trapping adaptations is against the finding of Brown and Khamphoukeo (2010), where trapping rodents is the most effective and important rodent control strategy. In fact, rodents are not easy to trap and may show trapping avoidances. This problem is largely associated with the ability of rodents to sense human smells on the traps, and inappropriate trapping procedures (setting the trap defectively, using inadequate and low-quality baiting food, and placing traps closely) (Hill, 2008). This suggests the need for a community wide education and training program on trap related problems and possible solutions.

The farmers in the study area used domestic cats as a natural enemy to control rodent pests in storage facilities. Owning local cats in the residential house is a widely employed biological rodent control method in different parts of Ethiopia (Yonas *et al*., 2010; 2014; Tomass *et al*., 2020). This practice is well documented and has a major suppressive effect on the local rat population (Hill, 2008). The effectiveness of this rodent management strategy in the area contravenes from other findings from other parts of Ethiopia (Kasso, 2013; Yonas *et al*., 2014) and elsewhere (Brown and Khamphoukeo, 2010). The possible explanation to this state of affairs could be due to the cost effectiveness of this method, and the differences in familiarity and availability of other rodent control measures between these areas. Such favor in the study area might be associated with farmers’ rodenticides and trapping adaptation claims, and the fear of the side effects of rodenticides. Kasso (2013) has argued that the use of domestic cats to control rodent pests is not equally effective in all areas because cats avoid catching and consuming rodents. Little is known about use of the field sanitation in rodent pest management among the farmers.

## Conclusion

Rodents cause considerable damage to agricultural crops and properties of small farm holders of Wenchi highlands. The farmers perceived rodents as a pest to their crops and created a nuisance to them. They declined to manage rodent damages in the crop fields. Owning cats, rodent trapping and using rodenticides were employed as indoor management strategies. The outcomes of this study suggest that rodent pests are a threat to food security in the area. Conducting on-field damage assessment and community level education programs are critical to estimate the actual damage rodents inflict in the field and to awaken the farmers for rodent management.

## Acknowledgements

The authors are grateful to Wolkite University and Addis Ababa University for their financial and material support. We are also indebted to acknowledge our field assistant, Malela Shiferaw for his aid during data collection, and our study participants for their willingness to participate in the study.

